# RelQ-mediated alarmone signaling regulates growth, sporulation, and stress-induced biofilm formation in *Clostridioides difficile*

**DOI:** 10.1101/2024.02.14.580318

**Authors:** Areej Malik, Adenrele Oludiran, Asia Poudel, Orlando Berumen Alvarez, Charles Woodward, Erin B. Purcell

**Affiliations:** Biomedical Sciences Program, Old Dominion University, Norfolk, Virginia, 23529, USA; Department of Chemistry and Biochemistry, Old Dominion University, Norfolk, Virginia, 23529, USA

## Abstract

The bacterial stringent response (SR) is a conserved transcriptional reprogramming pathway mediated by the nucleotide signaling alarmones, (pp)pGpp. The SR has been implicated in antibiotic survival in *Clostridioides difficile*, a biofilm- and spore-forming pathogen that causes resilient, highly recurrent *C. difficile* infections. The role of the SR in other processes and the effectors by which it regulates *C. difficile* physiology are unknown. *C. difficile* RelQ is a clostridial alarmone synthetase. Deletion of *relQ* dysregulates *C. difficile* growth in unstressed conditions, affects susceptibility to antibiotic and oxidative stressors, and drastically reduces biofilm formation. While wild-type *C. difficile* displays increased biofilm formation in the presence of sub-lethal stress, the Δ*relQ* strain cannot upregulate biofilm production in response to stress. Deletion of *relQ* slows spore accumulation in planktonic cultures but accelerates it in biofilms. This work establishes biofilm formation and sporulation as alarmone-mediated processes in *C. difficile* and reveals the importance of RelQ in stress-induced biofilm regulation.

## INTRODUCTION

*Clostridioides difficile* (*C. difficile*) is a Gram-positive, biofilm- and spore-forming obligate anaerobic pathogen that causes *Clostridioides difficile* infection (CDI) (1). This pathogen colonizes the mammalian large intestine after the disruption of beneficial commensal bacteria due to antibiotic treatment or advanced age (2). Antibiotic treatment for CDI further exacerbates the dysbiosis of the intestinal microbiota, which results in the recurrence of CDI (3). The chance of recurrent infection increases from 30 % to 45 % to 64 % after each successive antibiotic treatment for CDI, a phenomenon known as the ‘recurrence elevator’ (3). In the United States, *C. difficile* is responsible for approximately 500,000 infections, 25,000 deaths, and roughly $6 billion in hospitalization costs every year (4). Patients with severe or recurring CDIs are commonly treated with the antibiotics fidaxomicin, vancomycin, or metronidazole (5).

The bacterial stringent response (SR) is a conserved transcriptional regulatory mechanism that is activated when bacteria are exposed to stressful conditions. It allows bacteria to rapidly adjust to stress by temporarily inhibiting growth and cell division while simultaneously prompting the transcription of genes involved in stress survival (6, 7). In diverse microbes, the bacterial stringent response may be involved in tolerance of antibiotics, survival against antimicrobial peptides, inhibiting growth, resistance to oxidative stress, expression of genes governing virulence traits such as biofilm formation and sporulation, and/or competitive advantage against other microbes (8).

The SR is controlled by the accumulation of intracellular alarmone nucleotides that are collectively known as (pp)pGpp (9, 10). These alarmones are metabolized by bifunctional synthetase/hydrolase enzymes from the RSH or Rel (RelA/SpoT homology) family, monofunctional small alarmone hydrolases (SAH), and monofunctional small alarmone synthetases (SAS) (11). The SR is induced when bacteria are exposed to extracellular stresses that can include but are not limited to nutrient deprivation, antibiotic treatment, or oxidative stress (6, 12, 13). Firmicutes species generally encode only one RSH but can have one or two SASs, typically from the RelP and RelQ families (11). The highly conserved synthetase domains of RSH and SAS enzymes transfer a pyrophosphate from ATP to the 3’ OH of guanosine nucleotide precursors. Typically, the utilization of GMP, GDP, or GTP results in the synthesis of pGpp, ppGpp, or pppGpp, respectively (9, 10). *C. difficile* encodes highly conserved RSH and RelQ homologs whose alarmone synthesis contributes to antibiotic survival (10, 14, 15). However, it was recently discovered that both C. *difficile* synthetases must hydrolyze a 5’ phosphate or pyrophosphate from their GDP or GTP substrates while transferring the ATP-derived 3’ pyrophosphate, leaving behind a 5’ monophosphate and resulting in exclusive synthesis of pGpp (10, 14, 15).

Biofilm formation can be triggered by extracellular stresses. The process starts off with planktonic (free-floating) cells setting and attaching onto solid surfaces in single-species or polymicrobial communities (16). Biofilms are multilayered and have a protective matrix surrounding the cells, which is made up of bacterial proteins, extracellular DNA, and polysaccharides and which protects the cells from antibiotics and host immune responses (17, 18). Compared to free-living cells, cells within biofilms also undergo physiological changes such as decreased growth rate and altered transcriptional activity (19). The ability of *Clostridioides difficile* to form biofilms is one of the reasons that the pathogen can survive antibiotic treatment or exposure to immune stresses *in vivo* (17, 20, 21). The structure and composition of biofilm is dependent on the strain and biofilm age (17, 18). Some *C. difficile* strains are able to form more robust biofilms *in vivo*, resulting in higher antibiotic resistance (22). *Clostridioides difficile* strains isolated from patients with recurring CDIs produce more biofilms than strains isolated from patients whose CDIs resolved after one round of antibiotic treatment (23). This suggests *C. difficile* biofilms serve as a reservoir for recurrent infections (23, 24). It has also been shown that sublethal concentrations of the antibiotics vancomycin and metronidazole stimulate biofilm formation and thereby reduce antibiotic susceptibility and increase persistence and recurrence of CDIs (17, 25, 26).

The ability to form spores is critical for dissemination of *C. difficile* into the surrounding environment and spread to new patients (27, 28). The protective bacterial spore structure has characteristics that are different than that of a vegetative cell, which aid in bacterial resistance and durability in the environment (29, 30). Many of the layers surrounding the spore core contribute to bacterial spore resistance (28, 30–35). The spore core is highly dehydrated and composed of DNA, enzymes, tRNA, and ribosomes (30). It is known that antibiotics do not eradicate spores (36).

While metabolically active vegetative *C. difficile* cells are obligate anaerobes and are killed within minutes by exposure to atmospheric oxygen, spores can survive desiccation and exposure to environmental oxygen for months (37). Within the intestinal tract of a susceptible host, *C. difficile* spores germinate after exposure to primary bile acids and the amino acids alanine and glycine (38). Beneficial commensal microbes limit *C. difficile* germination and colonization by depleting nutrients such as amino acids and converting primary bile acids to secondary bile acids, which inhibit *C. difficile* growth, but these protective factors are diminished when advanced age or antibiotic treatment results in a disruption in the intestinal microbiota (39, 40).

While biofilm formation and sporulation are divergent developmental fates for an individual cell, they are complementary processes for a population. The majority of cells in 3-day old *C. difficile* biofilms are vegetative cells, but spores outnumber vegetative cells in 6-day old biofilms (18, 21). These results indicate that biofilms can serve as reservoirs for spores, thus increasing bacterial resistance to a variety of treatments (24, 41). While both processes are important for disease persistence, recurrence, and transmission, neither is completely understood in *C. difficile.* In spite of the high clinical significance of *Clostridioides difficile* infection, the signaling pathway that results in the transition of a dormant spore to a metabolically active vegetative cell is not completely understood in *C. difficile* (42). Some regulators that affect sporulation have been identified but lack conservation with other sporulating bacteria, and it is still not known how these regulators respond to external conditions (43). Similarly, it is known that biofilm formation is affected by the nucleotide signals c-di-GMP and c-di-AMP in *C. difficile* but the extracellular signals that trigger their synthesis and degradation are mostly unknown (44, 45).

The stringent response (SR) is conserved across almost all bacterial species, but the specific activating signals, effectors and resulting phenotypes governed by the SR vary between species (8, 46). The SR has been linked to virulence and biofilm formation in multiple bacterial species including *Streptococcus mutans*, *Pseudomonas aeruginosa*, *Enterococcus faecalis*, *Vibrio cholerae*, *Bordetella pertussis, and Escherichia coli* (47–53). The SR also plays a significant role in tolerance of *P. aeruginosa* and *S. aureus* biofilms to antibiotics and may stimulate biofilm formation in *Staphylococcus aureus* (48, 54–57). The SR also affects sporulation in *Bacillus subtilis,* and antibiotic tolerance in *Clostridioides difficile* (15, 58). Antibiotic exposure, oxidative stress, and metal starvation have been shown to upregulate transcription of clostridial synthetase genes, with *rsh* being more transcriptionally responsive than *relQ* (14, 15). Here we deleted the *relQ* gene in *C. difficile* 630Δ*erm* and observed that the knockout strain has reduced biofilm formation in all the conditions that were studied and has lost the ability to upregulate biofilm formation in response to stress. We further report that *relQ* deletion delays sporulation in liquid cultures and stimulates spore accumulation in biofilms. While RelQ is not the only clostridial alarmone synthetase, it does have a readily discernible role regulating stress-responsive biofilm formation and sporulation in *Clostridioides difficile*.

## RESULTS

### *relQ* regulates *Clostridioides difficile* 630Δ*erm* growth

Allele-coupled exchange (ACE) was used to excise the *relQ* open reading frame from the *C. difficile* 630Δ*erm* genome (Fig. S1). Growth of *C. difficile* 630Δ*erm* (wild-type) and 630Δ*erm* Δ*relQ* (knockout) strains was compared in BHIS broth. The duration of the lag phase, before exponential growth becomes visible, is similar for wild-type and knockout strains (Fig. 1). Once growth begins, the strains rapidly diverge. The wild-type strain exhibits ∼ 4 h of exponential growth with a doubling time of 1.5 h, followed by a transition into stationary phase at an optical density at 600 nm (OD_600_) of 0.55. The OD_600_ remains stable at 0.5 for the remainder of the 24 h experiment (Fig. 1). By contrast, the knockout strain has a doubling time of 0.79 h, until it reaches an OD_600_ of 0.95, followed by a continuous decline through the rest of the 24 h experiment (Fig. 1). Within 16 h, the knockout cell density has dropped below that of the wild-type (Fig. 1).

**FIG 1.**
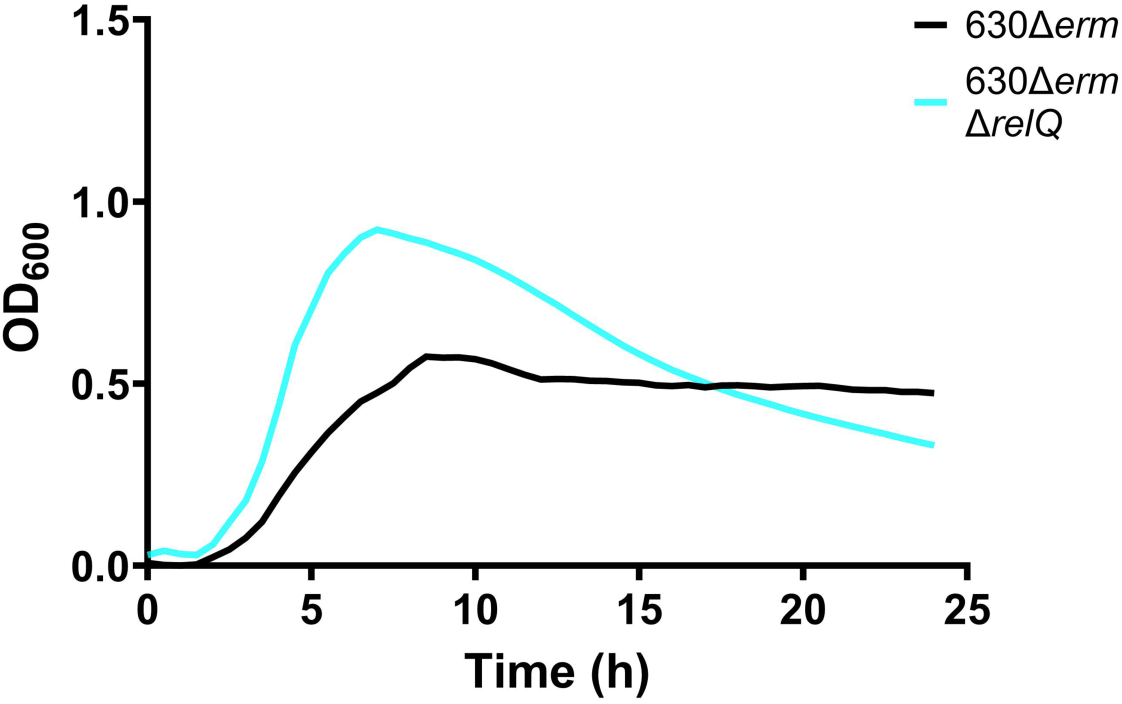
*Clostridioides difficile* 630Δ*erm* and 630Δ*erm* Δ*relQ* growth in BHIS broth. Growth in a 96-well plate of wild-type and knockout cultures in BHIS broth over 24 h. Data represents the mean of at least eight independent biologicals measured as duplicates.

### *relQ* deletion has non-uniform effects on *C. difficile* 630Δ*erm* growth during stress

We have previously demonstrated that the activity of the *relQ* promoter in *C. difficile* strains 630Δ*erm* and R20291 is stimulated by 2 h of exposure to sub-inhibitory concentrations of antibiotics in a strain-specific manner (14, 15). The *relQ* promoter was responsive to fidaxomicin in R20291 and to vancomycin in 630Δ*erm* but did not respond to metronidazole in either strain. (14, 15). To assess the impact of *relQ* on the cellular response to non-lethal antibiotic stress, we compared growth of the wild-type and knockout strains in the presence of antibiotics.

We found that while the wild-type and knockout strains exhibit different growth kinetics, both are delayed for several hours by the initial presence of a single dose of fidaxomicin (Fig. 2A and 2B). The knockout strain is slightly more sensitive to the antibiotic, exhibiting growth delays at concentrations of 0.01 µg/mL and above, while the wild-type strain shows no growth delay below 0.02 µg/mL (Fig. 2A and 2B). Similarly, 0.08 µg/mL fidaxomicin, the highest concentration tested, delayed wild-type growth by 14 h and knockout growth by 18 h (Fig. 2A and 2B). The wild-type and knockout strains both showed a negligible response to 0.25, 0.5, and 1.0 µg/mL vancomycin concentrations (Fig. 2C and 2D). The highest tested vancomycin concentration, 2.0 µg/mL, meaningfully suppressed wild-type growth for 24 h but had no apparent effect on knockout growth (Fig. 2C and 2D). The wild-type strain showed no negligible response to all the metronidazole concentrations that were tested (Fig. 2E). Unlike the wild-type strain, the knockout strain was slightly more sensitive to metronidazole, showing a modest growth defect at the concentration of 0.30 µg/mL (Fig. 2F).

**FIG 2.**
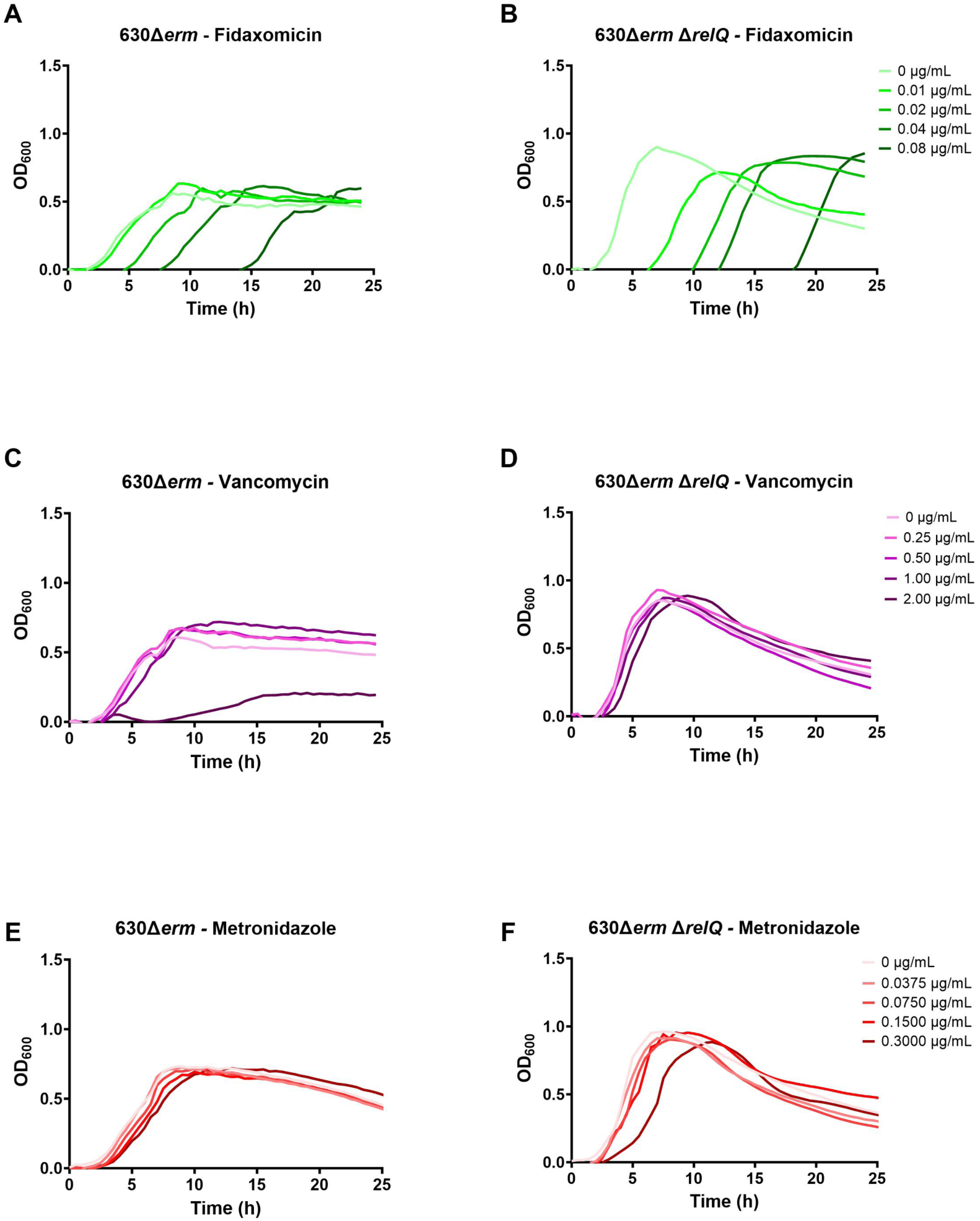
Effect of sublethal antibiotics on growth of *C. difficile* 630Δ*erm* and 630Δ*erm* Δ*relQ* strains. Growth of wild-type and knockout cultures in the presence of indicated concentrations of (A and B) fidaxomicin, (C and D) vancomycin, and (E and F) metronidazole. Data represents the mean of three independent biologicals measured as duplicates.

As an anaerobe, *C. difficile* is vulnerable to oxidative stress from molecular oxygen and transition metals (59, 60). Copper sulfate, a transition metal, can be used to create oxidative stress in an anaerobic environment (60). In the past our lab has demonstrated that 4.0 µM copper sulfate inhibits *C. difficile* 630Δ*erm* growth, although its effect on *rsh* and *relQ* transcription has not been assessed (60). Diamide simulates exposure to oxidative stress by initiating the formation of disulfide bonds in proteins (61). We have previously shown that 3 h treatment with 1.0 mM diamide activated the clostridial *rsh* promoter, although other stressors such as copper sulfate did not affect transcription of clostridial synthetases. (14). Here we tested the effect of sub-inhibitory exposure to copper sulfate and diamide on the growth of the wild-type and knockout strains. Neither oxidant affected wild-type growth (Fig. 3A and 3C). A slight decrease in growth of the knockout strain was observed when treated with 0.50 µM copper sulfate or 1.0 mM diamide (Fig. 3B and 3D). The absence of *relQ* disrupts the regulation of *C. difficile* 630Δ*erm* growth under normal conditions. Nevertheless, the impact of *relQ* deletion on cellular responses to sublethal stresses examined in this study was varied. In the absence of *relQ*, *C. difficile* 630Δ*erm* displays increased susceptibility to fidaxomicin and less susceptibility to vancomycin. However, *relQ* deletion does not meaningfully affect the growth of *C. difficile* 630Δ*erm* in the presence of metronidazole or oxidative stress.

**FIG 3.**
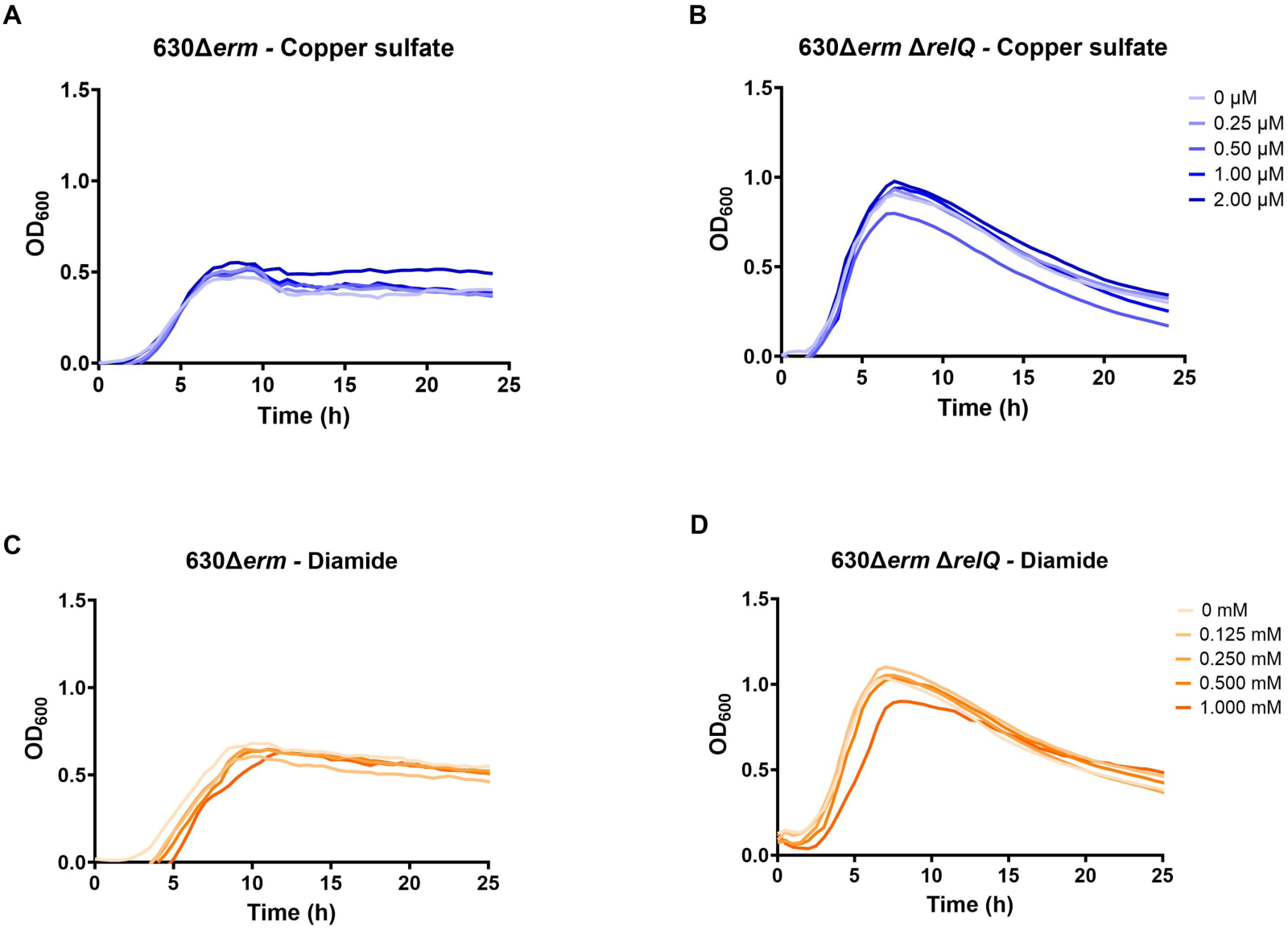
Effect of sublethal oxidants on growth of *C. difficile* 630Δ*erm* and 630Δ*erm* Δ*relQ* strains. Growth of wild-type and knockout cultures in the presence of indicated concentrations of (A and B) copper sulfate and (C and D) diamide. Data represents the mean of three independent biologicals measured as duplicates.

### *relQ* regulates *C. difficile* 630Δ*erm* stress-responsive biofilm formation

To determine whether RelQ-mediated alarmone signaling affects biofilm production in *C. difficile* 630*Δerm*, biofilms were grown in BHIS broth for 16, 24, and 48 h. After 16 h, no significant difference in biofilm formation was observed between the strains (Fig. 4). The wild-type strain showed substantially increased biofilm formation at 24 h, but the knockout did not (Fig. 4). Biofilm formation by both strains decreased between 24 and 48 h but remained much higher for the wild-type strain (Fig. 4).

**FIG 4.**
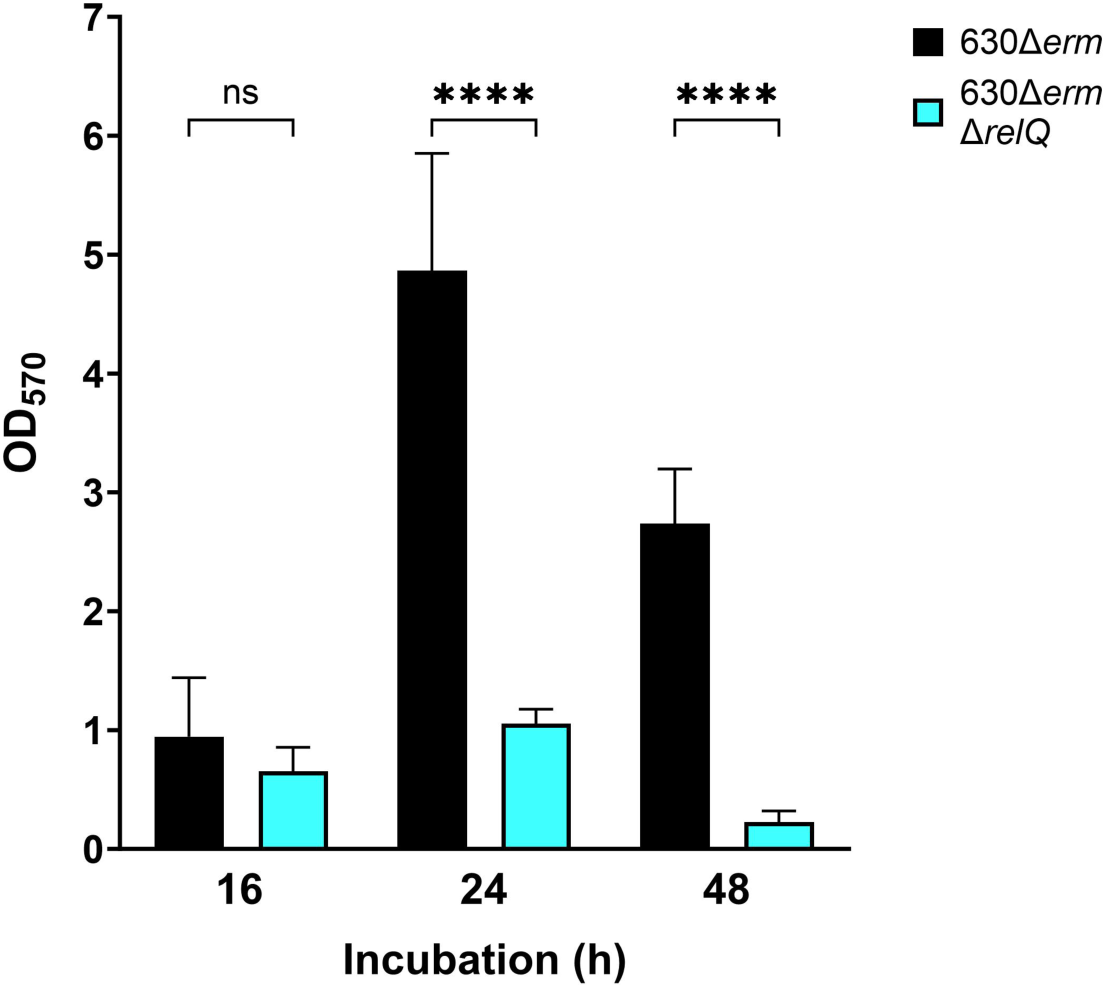
Biofilm production by *C. difficile* 630Δ*erm* and 630Δ*erm* Δ*relQ* strains. Biofilms were grown for 16 h, 24 h or 48 h before washing and staining with crystal violet. Data represents the mean and standard deviation of at least four independent biologicals measured in duplicate. Wild-type and knockout measurements at each time point were compared by one-way analysis of variance (ANOVA). *\*\*\*\*, p* < 0.0001; ns, not significant.

It has previously been shown that sublethal concentrations of antibiotics, including metronidazole and vancomycin, stimulate *C. difficile* biofilm formation (17, 25, 26). Our next step was to determine if *relQ* deletion affected the upregulation of biofilm formation as a response to specific stresses.

We initially tested biofilm formation in media containing 0.01 µg/mL fidaxomicin, 1.0 µg/mL vancomycin, and 0.3 µg/mL metronidazole. These were the highest antibiotic concentrations that allowed for growth of the wild-type strain (Fig. 2). The concentration of 1.0 µg/mL vancomycin did not show any noticeable effect on biofilm formation for both strains (data not shown). On the contrary, the presence of 0.3 µg/mL metronidazole did not stimulate biofilm formation for the wild-type strain but inhibited biofilm production for the knockout strain. (data not shown). We subsequently tested sub-inhibitory concentrations of 0.01 µg/mL fidaxomicin, 0.5 µg/mL vancomycin, and 0.08 µg/mL metronidazole and found that all these conditions stimulated a significant increase in 24 h biofilm formation by wild-type strain (Fig. 5). Contrary to the wild-type strain results, when the knockout strain was exposed to fidaxomicin, a significant decrease in biofilm formation was observed (Fig. 5A). Biofilm formation by the knockout strain showed no response to vancomycin or metronidazole stress, in contrast to the increases observed by the wild-type strain (Fig. 5B and 5C).

**FIG 5.**
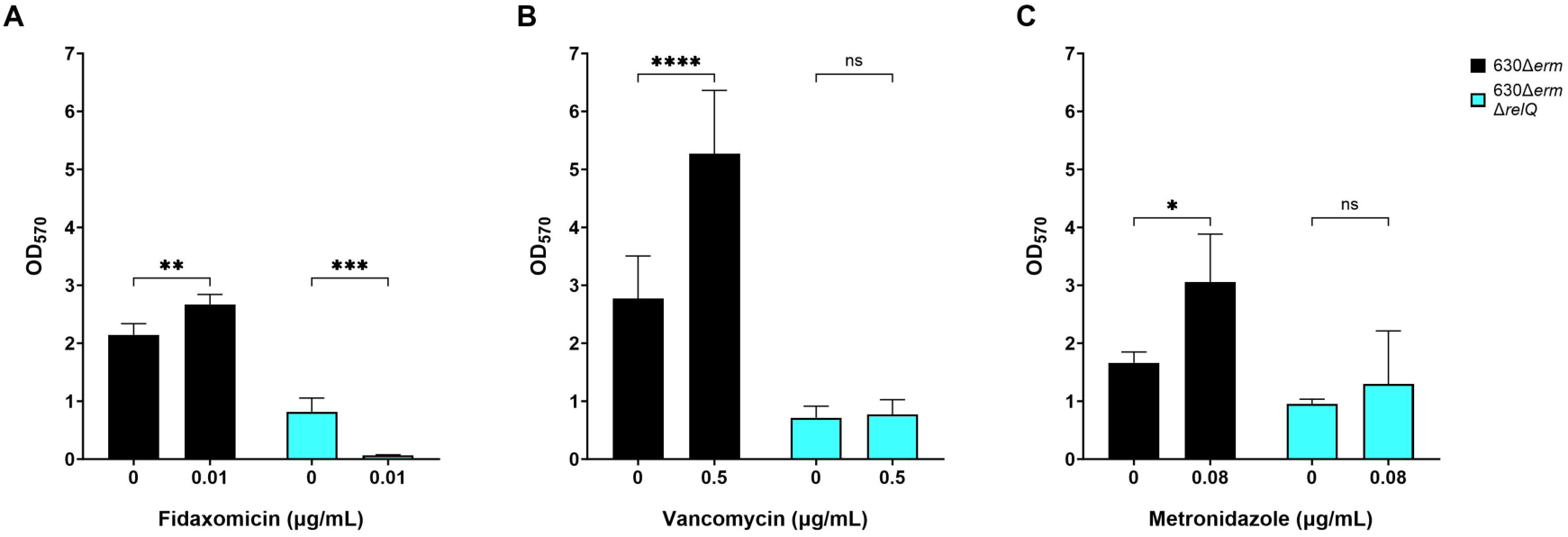
Effect of sublethal antibiotics on biofilm production by *C. difficile* 630Δ*erm* and 630Δ*erm* Δ*relQ* strains. Biofilms for both strains were grown for 24 h in the presence and absence of (A) 0.01 µg/mL fidaxomicin, (B) 0.5 µg/mL vancomycin, or (C) 0.08 µg/mL metronidazole before washing and staining with crystal violet. Data represents the mean and standard deviation of at least four independent biologicals measured in duplicate. Conditions with antibiotics were compared to untreated controls by one-way analysis of variance (ANOVA). ****, *p* < 0.0001*; ***, p* < 0.0005*; **, p* < 0.01*; *, p* < 0.05; ns, not significant.

Similar results were observed for biofilm formation when both strains were subjected to sub-lethal oxidative stress (Fig. 6). Treatment with 2.0 µM copper sulfate had no discernible impact on biofilm production for the wild-type and knockout strains (data not shown). Decreasing the copper sulfate concentration to 1.0 µM resulted in a significantly higher amount of biofilm formation by the wild-type strain while the knockout strain had no significant biofilm response to the copper stress (Fig. 6A). Treatment with 0.125 mM diamide had no effect on the growth of either strain (Fig. 3C and 3D). This sublethal diamide concentration stimulated a significant amount of biofilm formation by the wild-type strain while the knockout strain was non-responsive (Fig. 6B).

**FIG 6.**
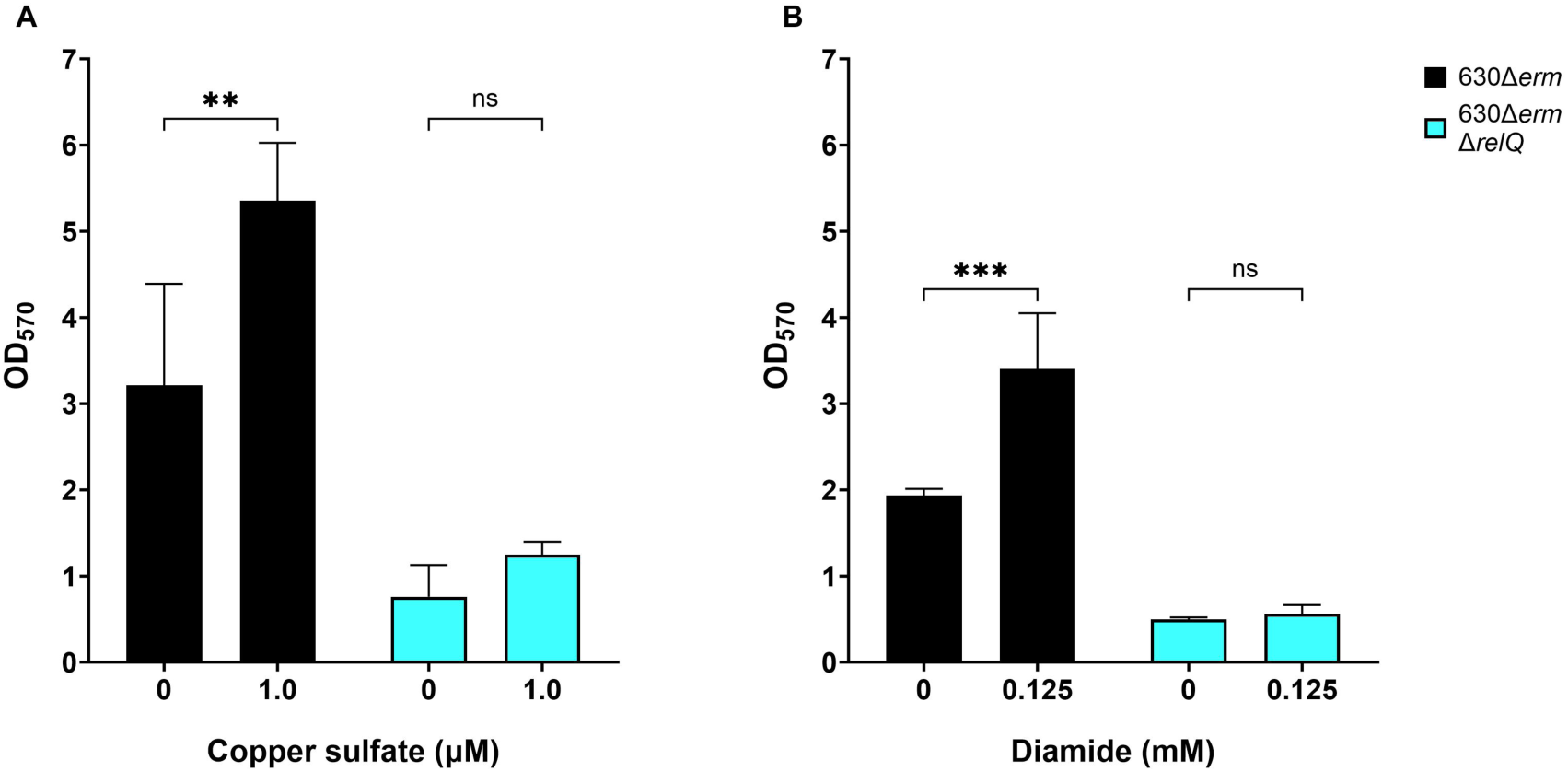
Effect of sublethal oxidants on biofilm production by *C. difficile* 630Δ*erm* and 630Δ*erm* Δ*relQ* strains. Biofilms were grown for 24 h in the presence and absence of (A) 1.0 µM copper sulfate and (B) 0.125 mM diamide. Data represents the mean and standard deviation of at least four independent biologicals measured in duplicate. Conditions with antibiotics were compared to untreated controls by one-way analysis of variance (ANOVA). ***, *p* < 0.0005; **, *p* < 0.01; ns, not significant.

### *relQ* deletion affects *C. difficile* spore accumulation in liquid cultures and biofilms

Sporulation and biofilm formation contribute to *C. difficile* virulence and recurrent infections. However, their coordinated regulation is poorly understood. We wanted to examine accumulation of spores in liquid cultures and in adherent biofilms. After 24 h, the wild-type forms very few spores in the liquid cultures (Fig. 7A). Between 24 and 48 h, an increase in number of spores is observed in liquid cultures (Fig. 7A). Wild-type spore accumulation in liquid culture does not increase substantially between 48 and 72 h (Fig. 7A). In liquid cultures, the knockout strain produces very few spores at 24 h or 48 h. However, an increase the number of viable knockout spores is observed between 48 and 72 h (Fig. 7A). In fact, the spore counts in liquid cultures after 72 h are similar between the two strains (Fig. 7A).

**FIG 7.**
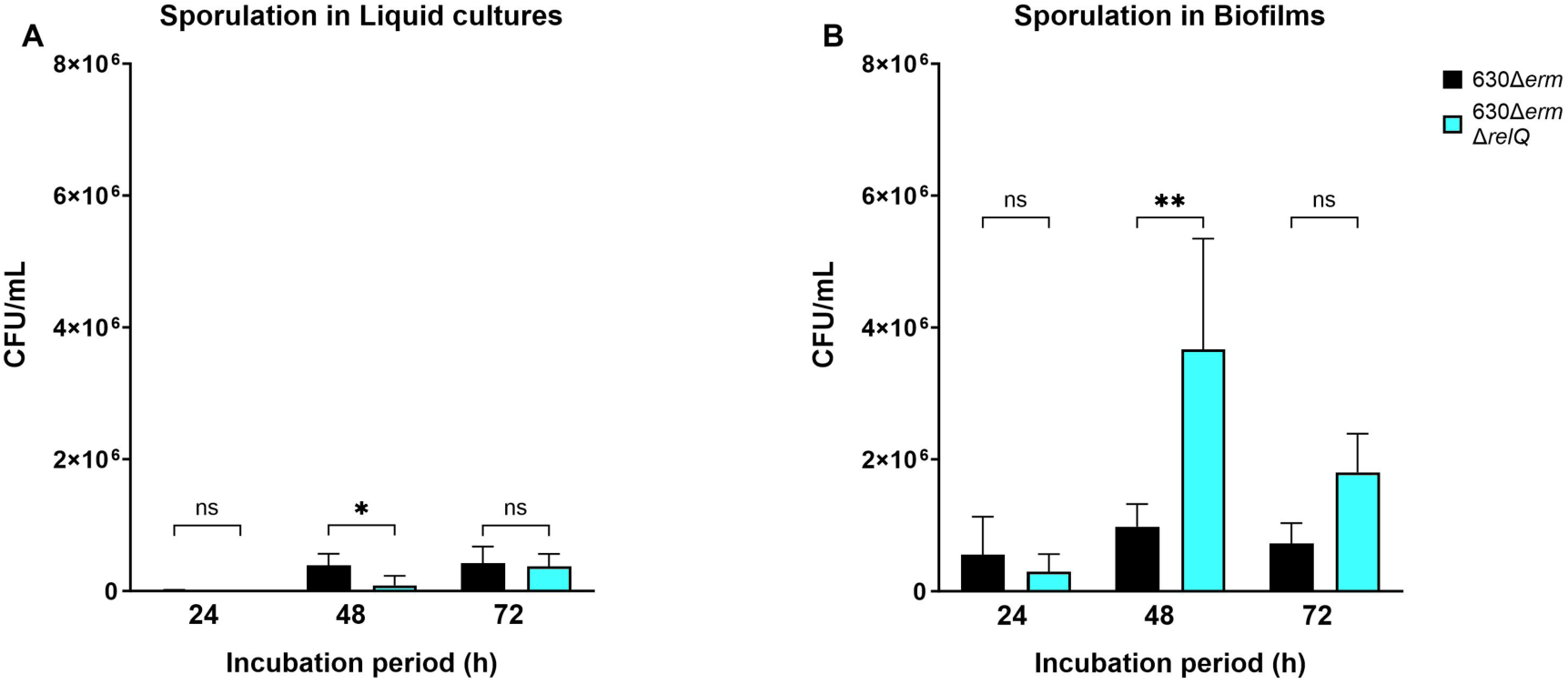
Spore production in liquid cultures or biofilms of *Clostridioides difficile* 630Δ*erm* and 630Δ*erm* Δ*relQ* strains. Spores of 630Δ*erm* and 630Δ*erm* Δ*relQ* were purified from (A) liquid broth or (B) washed biofilms after 24 h, 48 h, or 72 h of incubation. Data represents the mean and standard deviation of at least three independent biologicals measured in duplicate. Measurements were compared between the strains by one-way analysis of variance (ANOVA). *\*\*\*, p* < 0.0005; *\*\*, p* < 0.01 *\*, p* < 0.05; ns, not significant.

When the viable spores in washed biofilms were enumerated, there was no significant difference between the wild-type and knockout strains after 24 h, but the knockout strain showed significantly more spores in the 48 h biofilms (Fig. 7B). The spore concentration in the wild-type biofilms showed no significant difference between 48 and 72 h. In contrast, knockout biofilms exhibited reduced spore counts after 72 h, although the counts remained higher than those in the wild-type biofilms (Fig. 7B).

## DISCUSSION

The classical starvation-induced stringent response in Gram-positive Firmicutes is understood to be primarily mediated by a single RSH family enzyme, sometimes named Rel or RelA, while RelP and RelQ small alarmone synthetases more commonly play roles in basal alarmone synthesis and responses to cell wall stress (62–70). SAS have been linked to stress-induced biofilm formation in *S. aureus* (54). The drop in cytoplasmic GTP levels that results from alarmone synthesis has been linked to sporulation in *B. subtilis*, but to our knowledge alarmone-regulated sporulation in Firmicutes has not otherwise been demonstrated (58, 71).

Our results reveal that *relQ* deletion has meaningful and stress-specific effects on the exponential growth, biofilm formation, and spore accumulation of *C. difficile* 630Δ*erm* even in the presence of the intact bifunctional RSH. The absence of *relQ* disrupts the regulation of *C. difficile* 630Δ*erm* growth in unstressed conditions. However, the consequence of *relQ* deletion exhibit variability when the bacterium is exposed to sublethal stresses that were tested in this study. Without *relQ*, *C. difficile* 630Δ*erm* displays altered susceptibility to fidaxomicin and vancomycin but not to metronidazole or oxidants, underscoring the specificity of the RelQ-mediated response to individual extracellular stressors.

We have confirmed previous findings that exposure to sub-inhibitory antibiotic concentrations stimulated increased biofilm formation in wild-type *C. difficile* 630Δ*erm* and found that intact *relQ* is necessary for stress-induced biofilm upregulation. We also demonstrated that *C. difficile* increases biofilm accumulation in response to oxidative stress, which to our knowledge is the first report of this phenomenon and found that *relQ* is also necessary for oxidative stress-induced biofilm formation. As oxidative stress is employed by the innate immune system, this *relQ-*dependent biofilm stress response is likely to contribute to the high resilience of the pathogen within mammalian hosts (72).

*Clostridioides difficile* biofilms have high affinity for spores *in vitro* and have been proposed to serve as a reservoir for spores *in vivo*, suggesting that biofilm production and sporulation are complementary processes that are coordinately regulated (24). Deletion of the transcriptional regulator Spo0A abolishes sporulation in *C. difficile* R20291 and also results in notably diminished biofilm formation compared to the wild-type (21, 73). Biofilms formed by the *spo0A* mutant were prone to easy detachment from their growth surfaces and exhibited decreased resistance to oxygen stress (21). In our study, the *ΔrelQ* strain exhibited less biofilm formation, lost the ability to upregulate biofilm in response to stress, and altered the kinetics of spore accumulation in two different environments. The knockout accumulated spores more slowly in liquid broth and more rapidly in biofilms, despite having a lower total biofilm biomass than the wild-type at all timepoints. The loss of *relQ* resulted in a sparser biofilm containing a lower ratio of vegetative cells to associated spores, which could lead to altered persistence within a mammalian host. Future work with mixed-species biofilms and animal studies will be necessary to determine whether RelQ limits the pace of biofilm spore accumulation in more complex biofilms and whether the loss of this regulation is detrimental to survival and pathogenesis.

It is likely that we have observed evidence of two different roles played by RelQ in *C. difficile*. The inability of the knockout strain to upregulate biofilm formation in response to antibiotic or oxidant exposure indicates that RelQ contributes to the clostridial response to some extracellular stresses independently of RSH. However, in the absence of additional stress, the knockout strain exhibits accelerated exponential vegetative growth and reduced spore accumulation in liquid culture, suggesting that RelQ also restrains unstressed *C. difficile* growth and proliferation. Biofilms formed by the knockout strain are less substantial but contain more spores at earlier timepoints than those formed by the wild-type strain. Cell division and differentiation into either free-living planktonic cells, matrix-producing vegetative biofilm cells, or spores, are coordinately regulated, and it appears that and that pGpp produced by RelQ in response to growth conditions affects the distribution of the *C. difficile* population among these pathways. Future of examination of these processes in a pGpp-null (Δ*rsh*Δ*relQ*) strain could provide even more insight into the non-redundant functions of RSH and RelQ in this organism. It is clear that alarmone signaling affects many aspects of *C. difficile* physiology with potential relevance to infection persistence and recurrence.

## MATERIALS AND METHODS

### Bacterial strains and growth conditions

#Table S1 lists all the plasmids and bacterial strains that were used in this study. Table S2 lists all the primers that were used in this study. *Clostridioides difficile* strains were grown at 37 °C in brain heart infusion (BHI; VWR) medium supplemented with 5.0 % yeast extract (VWR) (74). These strains were cultivated in a Coy anaerobic chamber (Coy Laboratory) with an atmosphere of 10.0 % CO_2_, 5.0 % H_2_, and 85.0 % N_2_. *Escherichia coli* strains were grown in Luria-Bertani (LB; ThermoFisher Scientific) medium at 37 °C in a MaxQ 6000 incubator (ThermoFisher Scientific). Bacterial strains that carried plasmids were maintained using the following antibiotics at the indicated concentrations: 4 µg/mL or 50 µg/mL ampicillin (Amp) (Thomas Scientific), 100 ng/mL anhydrotetracycline (ATc) (Thomas Scientific), 15 µg/mL chloramphenicol (Cm) (Thomas Scientific), 10 µg/mL thiamphenicol (Tm) (ThermoFisher Scientific), 100 µg/mL kanamycin (Kan) (Thomas Scientific). All plastic consumables were brought into the anaerobic chamber for equilibration a minimum of 72 h before use.

### Plasmid and strain construction

The pMSR vector (provided by Olga Soutourina), containing the toxin gene CD2517.1 for counter-selection, was used for allele coupled exchange (ACE) in *C. difficile* 630Δ*erm* as described in reference (75). It was maintained in *E. coli* DH5α cells (EP109 in Table S1). NEBuilder HiFi DNA assembly was used to amplify the ‘RelQ arms,’ the upstream and downstream homology regions flanking *C. difficile* 630Δ*erm relQ* (CD630DERM_03450). The RelQ arms were amplified using primer pairs RelQ Arm 1_F + Arm 1_R and Arm 2_F + Arm 2_R (Table S2). The vector and amplified arms were digested using the PmeI cut site (NEB). The arms and vector were ligated using the Gibson Assembly cloning kit (NEB), thus generating pMSR::630Δ*erm*_*relQ*_arms, which was transformed into competent *E. coli* DH5α cells (EP111 in Table S1). The plasmid was confirmed by PCR using primer pair, RelQ Arm 1_F and Arm 2_R (Table S2). The pMSR::630Δ*erm*_*relQ*_arms vector was then transformed into competent *E. coli* HB101 cells carrying the pRK24 plasmid (EP113 in Table S1) to create the donor strain for conjugative transfer. The pRK24 plasmid is a derivative of a broad-host-range plasmid RP4 that is involved in the transfer of plasmid DNA into *C. difficile* strains (76).

A singly colony of the donor strain was inoculated in 3 mL of LB media supplemented with 15 µg/mL Cm and 50 µg/mL Amp. The culture was incubated for 14-16 h at 37 °C with continuous shaking at 250 rpm. The overnight culture was plated on LB agar plates supplemented with 15 µg/mL Cm and 50 µg/mL Amp. The plates were incubated overnight at 37 °C. Inside the anaerobic chamber, a single colony of the acceptor strain was inoculated in 3 mL of BHIS broth, which was then incubated at 37 °C for 14-16 hr. The lawns of *E. coli* donor cells were scraped from the agar plate and resuspended in 1 mL LB broth, centrifuged, and resuspended in 150 µL LB broth. Inside the anaerobic chamber, a 1:1 ratio of both donor and acceptor cell suspensions were mixed, and 50 µL spots were pipetted onto BHIS agar plates supplemented with 11 mM glucose (ThermoFisher Scientific) and 4 µg/mL Amp. The plates were incubated at 37 °C for 18 h inside the anaerobic chamber. After the 18 h incubation, the spots were scraped up with a sterile inoculating loop and suspended in 1 mL of BHIS broth before 100 µL of suspension was spread onto BHIS agar plates supplemented with 10 µg/mL Tm and 100 µg/mL Kan. These selective plates were incubated at 37 °C for 20-24 h. Colonies were then streaked twice onto fresh BHIS agar plates supplemented with 10 µg/mL Tm and 100 µg/mL Kan. The resulting colonies were then passaged onto BHIS agar plates supplemented with 10 µg/mL Tm to select for plasmid integration into the chromosome. The next step was to streak the colonies twice on BHIS agar plates supplemented with 100 ng/mL ATc to induce expression of the plasmid-encoded *CD2517.1* toxin and induce plasmid excision. These counter-selected colonies were then simultaneously streaked onto BHIS agar plates supplemented with 10 µg/mL Tm, and only colonies that had regained sensitivity to thiamphenicol were screened as potential knockouts. Potential knockouts were confirmed via PCR with primer pair RelQ Arm1_F and RelQ Arm 2_R (Table S2, Fig. S1).

### Growth curves

*C. difficile* 630Δ*erm* and 630Δ*erm* Δ*relQ* cultures were prepared by inoculating individual colonies into 3 mL of BHIS broth and grown for 14-16 h at 37 °C in the anaerobic chamber. The overnight cultures were diluted 1:20 into BHIS broth containing the indicated concentrations of fidaxomicin (Cayman Chemical), vancomycin (VWR), metronidazole (VWR), copper sulfate (Fisher Scientific), or diamide (MP Biomedicals) into sterile 96-well plates (Fisher Scientific). The plates were incubated at 37 °C for 24 h in a Stratus microplate reader (Stellar Scientific), which was set to record the optical density at 600 nm every 30 min.

### Biofilm formation, visualization, and quantification

Overnight cultures of *C. difficile* 630Δ*erm* and 630Δ*erm* Δ*relQ* were diluted 1:10 into BHIS broth containing the indicated concentrations of fidaxomicin, vancomycin, metronidazole, copper sulfate, or diamide. Biofilms were grown in a total volume of 2 mL in 24-well sterile non-treated plates (Corning) under anaerobic conditions at 37 °C for 24 h. After the incubation, the plates were removed from the chamber and the exteriors were sterilized with SporGon (Thomas Scientific), 10 % bleach, and 70 % ethanol (VWR). The supernatant was removed by pipetting and the adherent biofilms were washed with 1 mL of 1X phosphate buffered saline (PBS) solution. The washed biofilms were stained with 1 mL of 0.1 % crystal violet (Sigma Aldrich) in each well for 30 min at room temperature. After the 30 min incubation, the crystal violet was removed, and the stained biofilms were washed twice with 1X PBS. Stained biofilms were scraped up with sterile pipet tips and suspended in 1 mL of 70 % ethanol. A Bio-Tek synergy plate reader (Marshall Scientific) was used to record the optical density at 570 nm after shaking the plate at medium intensity.

### Viability assay of spores in liquid cultures

Individual colonies of *C. difficile* 630Δ*erm* and 630Δ*erm* Δ*relQ* strains were inoculated into 3 mL of BHIS broth. The cultures were incubated for 24, 48, or 72 h at 37 °C in a Coy anaerobic chamber. After the indicated incubation, duplicate 470 µL aliquots of each culture were removed from the chamber in 2 mL eppendorf tubes (VWR), which were sterilized with SporGon, 10 % bleach, and 70 % ethanol. The experimental tubes were opened and exposed to oxygen for 1 h and treated with 530 µL of 95 % ethanol, to get a final concentration of 50% ethanol, for 1 h with periodic mixing every 15 min, while the controls remained closed and were not treated with ethanol. After the 1 h treatment, the samples were brought back into the chamber and subsequently underwent serial dilution in BHIS broth. A 100 µL aliquot of the original and diluted samples were plated as duplicates on BHIS agar with and without 0.1% taurocholic acid. The plates were incubated at 37 °C for 48 h before the viable colonies were counted. The range for viable colony count was restricted between 20 to 350.

### Viability assays of spores in biofilms

Biofilms were inoculated, as described in “Biofilm formation, visualization, and quantification” methodology section, and incubated anaerobically at 37 °C for 24, 48, or 72 h. After the respective incubation, the plates were brought out of the chamber and the exteriors were sterilized. The supernatant was discarded and the adhered biofilms in each well were washed with 1 mL of 1X PBS. The biofilms were allowed to dry before being washed again with 1 mL of 1X PBS. The washed biofilms were scraped up in each well with a sterile pipet tip before being suspended in 1 mL of BHIS broth. The samples were taken into the chamber where they underwent a serial dilution in BHIS broth before being plated as duplicates on BHIS agar with and without 0.1% taurocholic acid. The plates were incubated at 37 °C for 48 h before the viable colonies were counted. The range for viable colony count was restricted between 20 to 300.

### Statistical analyses

The growth curve data are presented as the mean for each experiment. The biofilm and sporulation data are presented as the mean ± standard deviation. Prism 10 (GraphPad) was used for all statistical calculations, data fitting and plotting. Quantitative experiments were performed as duplicates and were repeated at least three times.

## ACKNOWLEDGEMENTS

We thank Olga Soutourina for the PMSR vector. This research was supported by NSF LEAPS-MPS 2213353 and by NIH 1 R15 AI156650-01. The ODU Biomedical Sciences program provided funding for supplies to Areej Malik.

## Notes

### Competing Interest Statement

The authors have declared no competing interest.

