## Supplemental Material for "RelQ-mediated alarmone signaling regulates growth, sporulation, and stress-induced biofilm formation in *Clostridioides difficile*"

**Table S1** Strains and plasmids used in this paper.

| Plasmids | | |
|---|---|---|
| **Name** | **Description** | **Reference** |
| pMSR | ACE vector for chromosomal manipulation of *C. difficile* 630 using a counter-selection marker (CD2517.1 toxin) | [1] |
| pRK24 | Used to facilitate in the transfer of plasmid DNA into *C. difficile* strains. | [2] |
| pMSR::630Δ*erm_relQ*_arms | Modified pMSR ACE vector with 630Δ*erm relQ* flanking sequences to target the *relQ* gene | This study |
| ***Escherichia coli*** | | |
| **Name** | **Description** | |
| DH5α | Gentoype: *fhuA2Δ(argF-lacZ)U169 phoA glnV44 Φ80Δ(lacZ)M15 gyrA96 recA1 relA1 endA1 thi-1 hsdR17* | NEB |
| EP109 | pMSR in DH5α | This study |
| EP111 | pMSR::630Δ*erm relQ* arms in DH5α | This study |
| RT270 | pRK24 in HB101 | This study |
| EP113 | pMSR::630Δ*erm_relQ*_arms and pRK24 in HB101 | This study |
| ***Clostridioides difficile*** | | |
| **Name** | **Description** | |
| 630Δ*erm* | CD630 strain lacking *erm*B | [3] |
| CEP46 | 630Δ*erm* lacking *relQ* | This study |

**Table S2.** Oligonucleotide primer sequences used in this paper.

| Cloning primers | | |
|---|---|---|
| **Name** | **Description and Sequence (5'-3')** | **Reference** |
| EP143 | RelQ Arm 1_fwd TTTTTTGTTACCCTAAGTTTGTTCAAGACCTAGGAGAG | This study |
| EP144 | RelQ Arm 1_rev GCCCCATTAAGCTCACCCTTTATTTGTTTTTATG | This study |
| EP145 | RelQ Arm 2_fwd AAGGGTGAGCTTAATGGGGCTGCCTCTTAATG | This study |
| EP146 | RelQ Arm 2_rev AGATTATCAAAAAGGAGTTTCCGAAATCCCTCAAATATTTATTTTTAATC | This study |

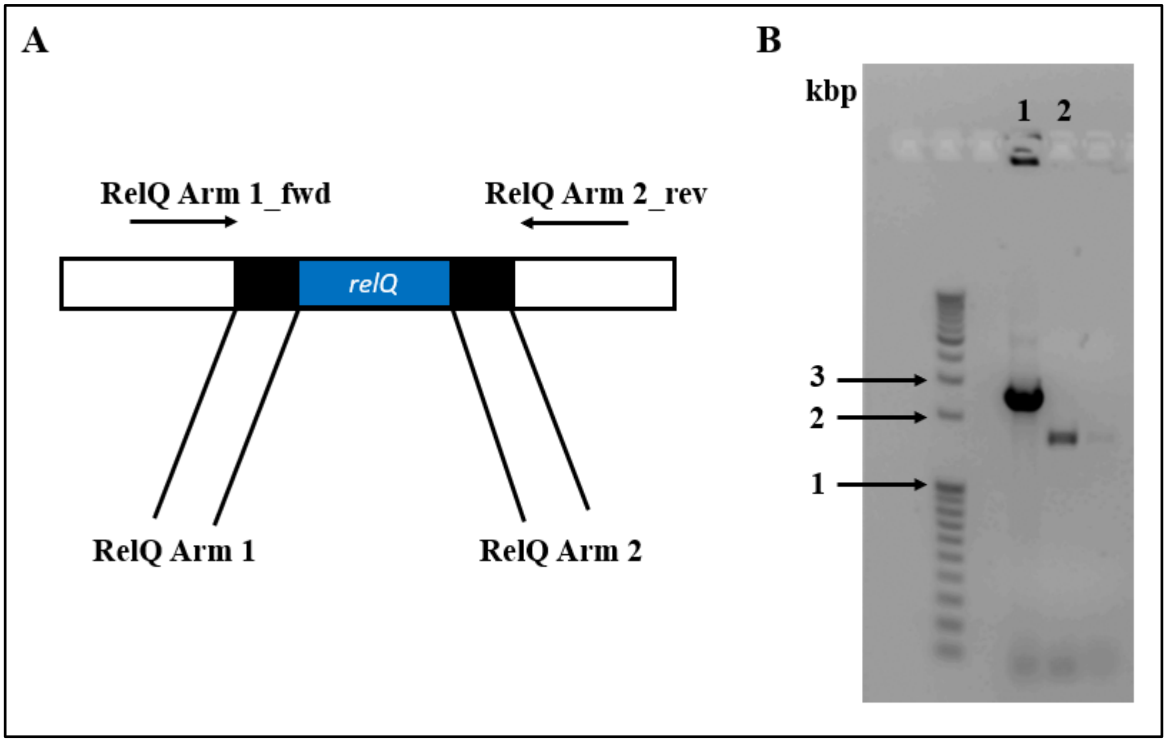


**FIG S1** Confirmation of presence and absence of *relQ* in *C. difficile* 630Δ*erm* and *C. difficile* 630Δ*erm* Δ*relQ* strains. (A) Pictured are the *relQ* open reading frame (blue) and the amplified flanking regions (black). The primers RelQ Arm1_fwd and RelQ Arm2_rev amplify a 2.5 kbp region from the wild-type genome. In the Δ*relQ* strain, only the flanking regions are present and these primers amplify a 1.7 kbp product. (B) The products amplified from *C. difficile* 630Δ*erm* (lane 1) and *C. difficile* 630Δ*erm* Δ*relQ* (lane 2) using RelQ Arm1_fwd and RelQ Arm2_rev are pictured on a 0.8% DNA agarose gel.
